# Transitions in interaction landscapes of multidrug combinations

**DOI:** 10.1101/367664

**Authors:** Tina Manzhu Kang, Bjørn Østman, Mauricio Cruz-Loya, Natalie Ann Lozano, Robert Damoiseaux, Van M. Savage, Pamela J. Yeh

## Abstract

Drug combinations are a promising strategy to increase killing efficiency and to decrease the likelihood of evolving resistance. A major challenge is to gain a detailed understanding of how drugs interact in a dose-specific manner, especially for interactions involving more than two drugs. Here we introduce a direct and intuitive visual representation that we term “interaction landscapes”. We use these landscapes to clearly show that the interaction type of two drugs typically transitions smoothly from antagonism to no interaction to synergy as drug doses increase. This finding contradicts prevailing assumptions that interaction type is always the same. Our results, from 56 interaction landscapes, are derived from all possible three-drug combinations among 8 antibiotics, each varied across a range of 7 concentrations and applied to a pathogenic *Escherichia coli* strain. Such comprehensive data and analysis are only recently possible through implementation of an automated high-throughput drug-delivery system and an explicit mathematical framework that disentangles pairwise versus three-way as well as net (any effect) versus emergent (requiring all three drugs) interactions. Altogether, these landscapes partly capture and encapsulate selective pressures that correspond to different dose regions and could help optimize treatment strategies. Consequently, interaction landscapes have profound consequences for choosing effective drug-dose combinations because there are regions where small changes in dose can cause large changes in pathogen killing efficiency and selective pressure.

## Introduction

Combination therapy is widely used to treat a number of chronic health issues such as cancer [1, 2], HIV [3, 4], hypertension [5] or multidrug resistant bacterial infections [6, 7]. Understanding the effects of these drug combinations and interactions among drugs is a major clinical concern and active research area [8–14]. A promising strategy for combatting the evolution of drug resistance is to use drugs in combination by effectively leveraging interactions. However, a detailed understanding of how three drugs interact in a dose-specific manner is challenging to examine and visualize. Gaining this understanding has importance both for devising optimal treatments and for leveraging selection pressure to combat evolution of resistance.

Measures for interactions are often evaluated based on a coarse-grained categorization of three interaction types: additive (no interaction), synergistic (combined effect greater than expected based on single-drug effects), and antagonistic (combined effect less than expected based on single-drug effects). Synergistic drug combinations, in which combining drugs enhances the effects of the individual drugs, are commonly prescribed for patients because they maximize efficacy at lower doses. However, previous work indicates that antagonism may be more beneficial for slowing down the rate of resistance evolution to the component drugs [10, 14], because it creates more complex or rugged fitness landscapes. Thus, simply knowing how interactions deviate from additivity towards synergy or antagonism is potentially a powerful indicator to anticipate effects of a specific drug combination on treatment and resistance development.

Nevertheless, in practice it becomes challenging to use this interaction categorization to optimize treatment strategy and leverage evolution of resistance due to the complexity of dose-dependent interactions. Many empirical studies of drug interaction are conducted at a fixed dose and thus can only measure a single interaction type for each specific drug combination (i.e., Bliss Independence) [10, 13, 15]. The Bliss independence model is one of the most commonly used measures of drug interactions because it is intuitive, simple to calculate, readily expandable to numerous interacting components, and experimentally less demanding because it only requires four measurements to classify a pairwise interaction.

When drug combinations do not have an unchanged interaction type with changing drug dose [16], there is a breakdown of common interaction definitions based on single-dose measurements. Several studies have now shown that changes in interactions based on doses is not just an abstract possibility but a reality for combinations of antibiotics, antifungal, and chemotherapeutic agents [15, 17–20]. More systematic studies are needed to find and understand general patterns and thus to avoid adverse effect that promote development of resistance and disease relapse. Such cases could occur when the interaction of a drug combination is defined at a specific dose combination and is extrapolated into a region of drug doses where the interaction is neither what is expected nor what is desirable. Until now, there have been no direct and intuitive visualizations of high-dimensional drug spaces that would help verify and more deeply understand the range of complexities in transitions among interaction types.

In this paper, we examine all possible three-drug combinations among 8 antibiotics, each varied across a range of 7 concentrations and applied to a pathogenic *Escherichia coli* strain. We introduce a new and direct visual representation of dose-dependent drug interactions that we term “interaction landscapes” (Figure 1). This approach is placed directly within the space of drug interactions where general inferences about consequences of interactions can be made quickly with extremely efficient use of the information in the data. Interestingly, because interactions are calculated from fitness differences, the interaction landscape is a visual representation that partly captures directions and strengths of selection pressures. Therefore, these interaction landscapes will help to analyze how drug-dose combinations affect treatment strategies, regions of positive or negative selection pressures, and evolution of resistance.

**Figure 1.**
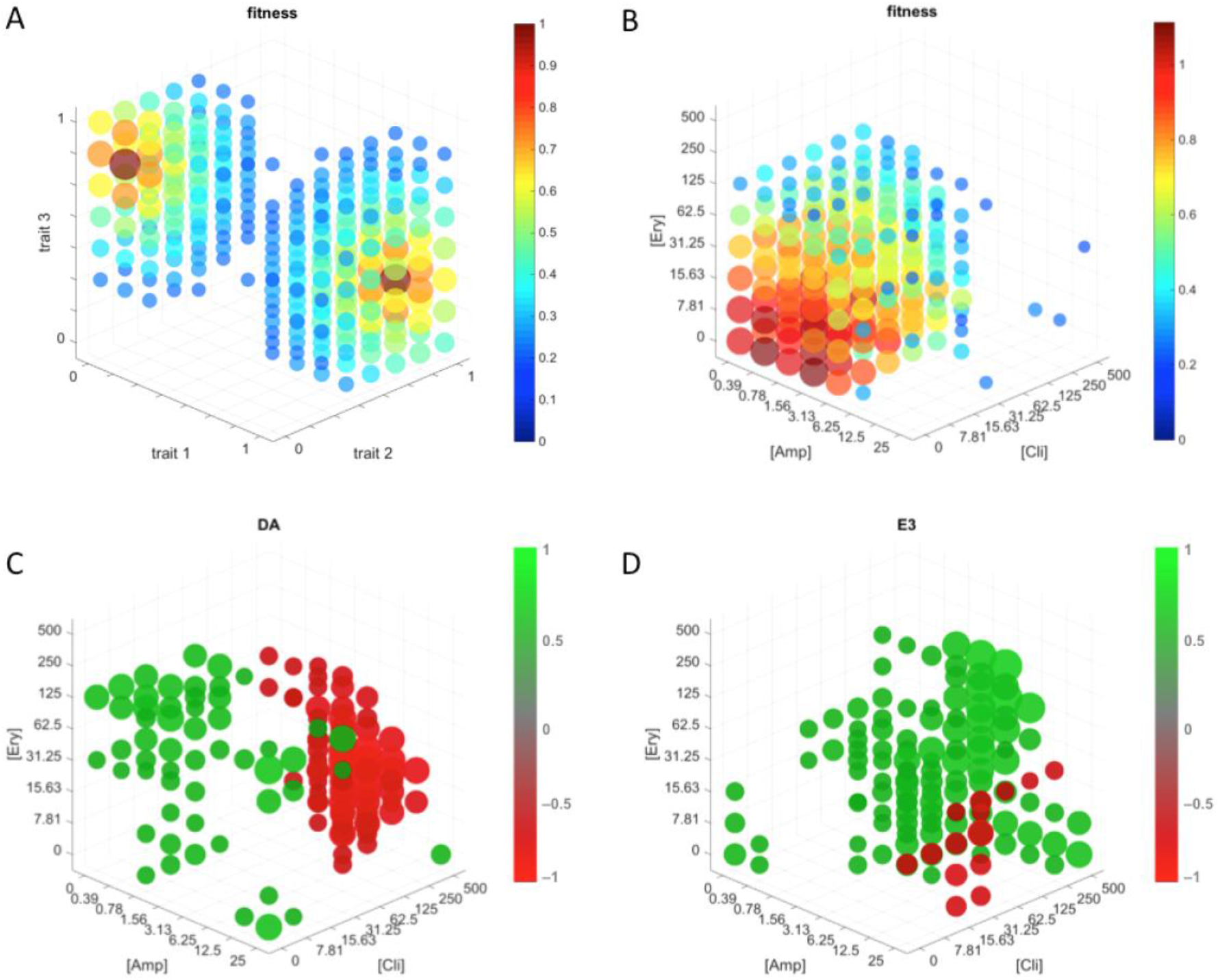
Overview and comparison of traditional fitness landscapes and interaction landscapes. (A) Schematic of a traditional phenotype-fitness landscape with fitness as a function of three continuous trait values. Two peaks are visible, showing that there are different combinations of trait values favored by selection. Values below 0.2 are not shown. (B) Fitness landscape with fitness as a function of drug concentration for AMP/CLI/ERY. Fitness is optimal when no drugs are administered. Values below 0.2 are not shown. (C) *DA* interaction landscape for AMP/CLI/ERY showing two distinct regimes where interactions are primarily antagonistic (green) and synergistic (red). Values between –0.5 and 0.5 are not shown. (D) E3 interaction landscape for AMP/CLI/ERY, showing how interactions can be dramatically different between net (DA) and emergent (E3) interactions.

Interaction landscapes, which are based on our high-throughput data and calculated from our mathematical framework, provide direct visualizations of local synergy or antagonism embedded within a larger interaction space and thus enable quantification and assessment of the directionality, pervasiveness, organization, and transition between regional synergy and antagonism. Consequently, we can use these landscapes to carefully investigate and answer the questions above. We expect broad implications of this general approach and ideas, including in environmental pollution and risk assessment of toxic chemical mixtures where the exposure is rarely a uniform dose.

## Results

Overall, we find that interaction types are often strongly dose-dependent, and that this is true for both lower-order (two drug) and higher-order (three drug) combinations. We typically observe smooth transitions between different interaction types and subspaces within a drug combination. Furthermore, net interactions tend to transition from antagonistic at low dose to synergy at high doses. For emergent interactions, higher doses often have the opposite effect and lead to more antagonism. These transitions happen quickly but smoothly. Finally, pairwise interactions can often be used to predict net three-drug interactions but not emergent three-drug interactions.

### Interaction type is dose dependent

Both lower-order (2-drug) and higher-order (3-drug) interactions are strongly dose dependent. To assess the effect of increasing dose on interaction in a two- drug case, we compared how sub-inhibitory concentrations of drug A interact with drug B at either a high dose or a low dose. In a three-drug combination, the interaction was examined with the third drug at a high and low dose. We measured interaction both at the overall net level (*DA*)—combined pairwise and three-way interactions—and at the emergent level (*E3*), where the pairwise interactions are subtracted from the net interactions so that only the truly three-way interaction part remains (Figure 2). The distributions of *DA* and *E3* among all combinations of drugs and doses are multimodal with peaks at synergy (*DA* = –1), additivity (*DA* = 0), and antagonism (*DA* = 1) (Fig. 3A and B). Smoothing the data results in a more continuous distribution (Fig. 3C and D). The peaks at the boundaries of synergy and antagonism were much less prominent (Fig. 3C and D), and low drug concentrations result primarily in net interactions that are additive or antagonistic (Fig. S1). Synergistic *DA* and *E3* interactions are mostly observed at intermediate and high concentrations with a dearth at low doses (Fig. 3C and D).

**Figure 2.**
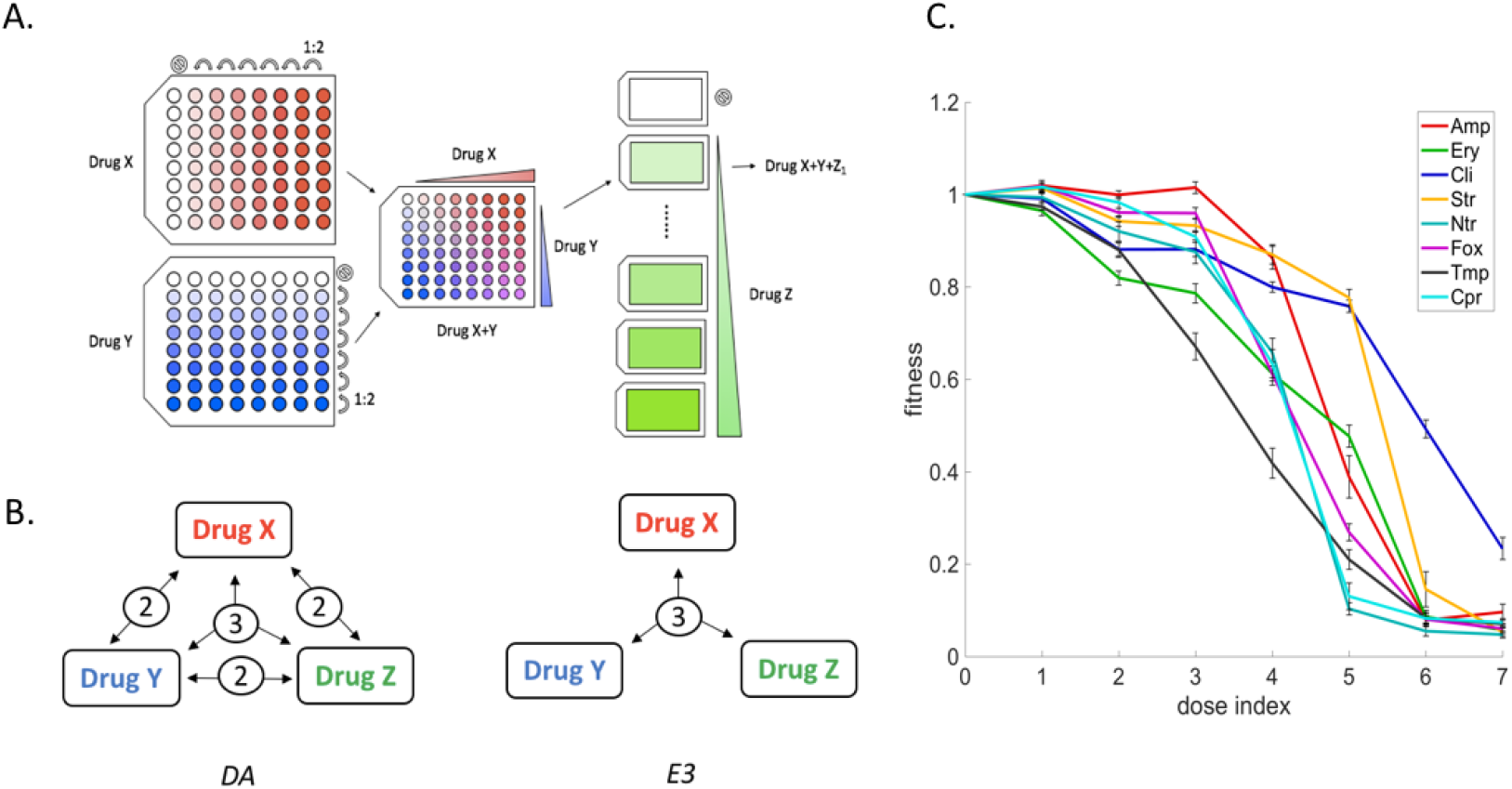
Schematic representation of experimental design. (A) For one triple-drug combination of X, Y, and Z, the drug X plate includes 7 steps of 2-fold serial dilutions (in red) plus no drug control (in white) going in the horizontal direction. Drug Y plate includes the same concentration gradient but in the vertical direction (in blue). Combining drug X and drug Y plates results in a 2-dimensional matrix of drug X+Y. Drug Z is composed of 7 plates each with one concentration across the full 7-drug gradient (in green). Each of the seven drug Z plates is transferred to a drug X+Y plate to form a matrix of X+Y+Z at one respective dose (drug X+Y+Z). Finally, a 3-dimensional matrix of all three drugs is constructed of all seven additions of Z into one plate of X+Y, plus a control where Z is zero. (B) For each drug-dose combination, the overall interaction of *DA* is calculated with three component pairwise interactions of drug X+Y, drug X+Z, and drug Y+Z (2), and the interaction when all three drug are present (3); while emergent three way *E3* represent the interaction of only at the three drug level. (C) Mean dose response curves for single drugs from the dilution scheme, where the dilution step is plotted as dose index and the fitness as a function of dose. AMP (20 replicates, red), ERY (21 replicates, green), CLI (21 replicates, blue), STR (21 replicates, orange), NTR (21 replicates, teal), FOX (20 replicates, purple), TMP (21 replicates, black), CPR (20 replicates, cyan).

**Figure 3.**
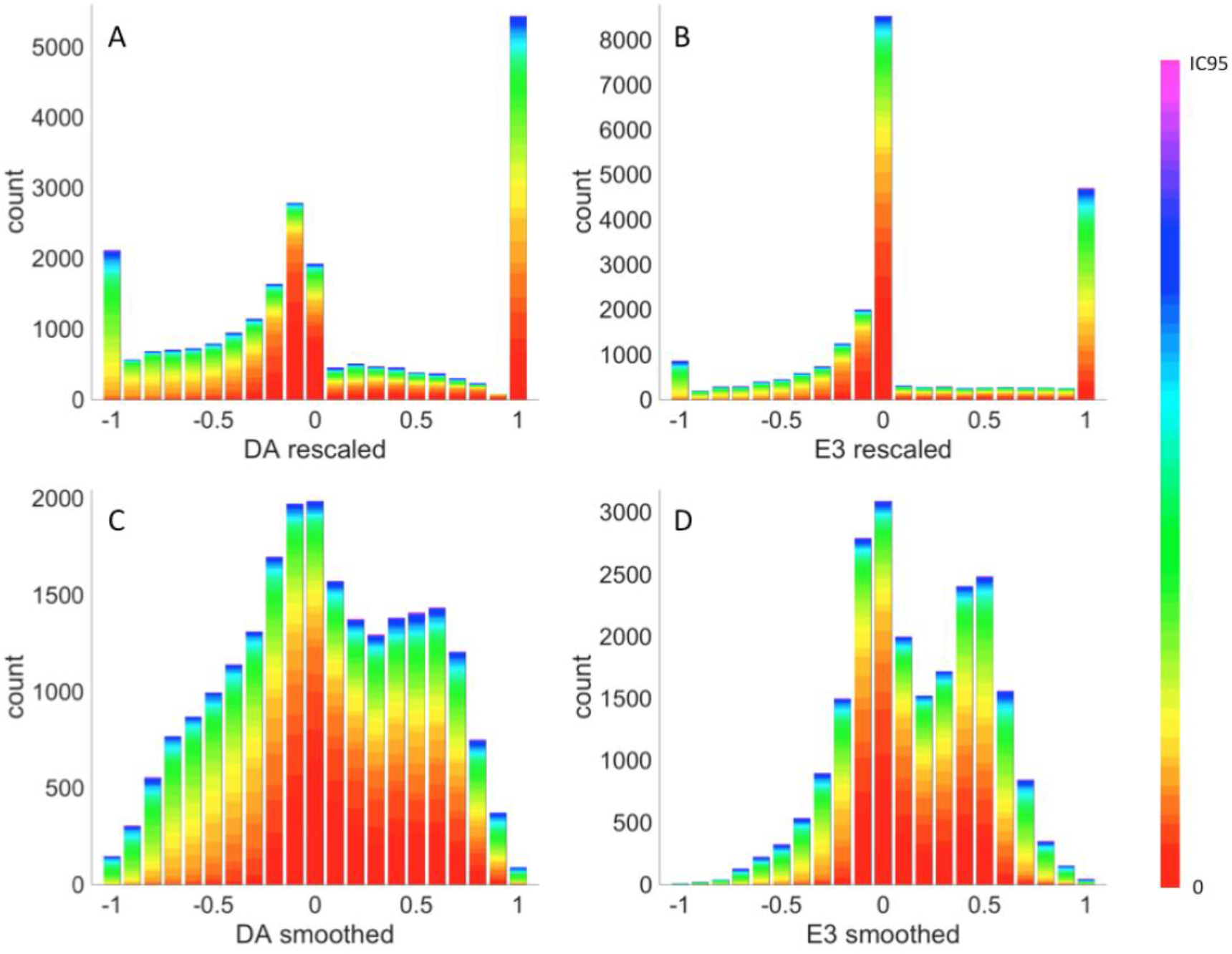
Rescaled net (*DA*) and emergent (*E3*) interaction distributions. Panel A and C show the overall net level (*DA*) which encompasses all component pairwise and three-way interactions. Panel B and D show interaction at the emergent level (*E3*), where the pairwise interactions are subtracted from the net interactions so that only the truly three-way interaction part remains. The colors correspond to drug concentration, where IC95 were used as the maxima (see methods). Drug concentrations above IC95 were counted as the maxima. (A) Rescaled *DA*, (B) rescaled *E3*, (C) smoothed *DA*, (D) smoothed *E3*. Low drug concentrations (red) result predominantly in additive (–0.5 < *DA* < 0.5) or antagonistic interactions (*DA* > 0.5). Higher concentrations (green to purple) are more evenly distributed among interaction types.

### Interaction type transitions

Interaction types tend to transition from antagonism and additivity at low doses to synergy at high doses for net three-way interaction, but to antagonism for emergent three-way interaction. For both *DA* and *E3*, the magnitude of the mean of all antagonistic interactions and the magnitude of the mean of all synergistic interactions each increase with the combined dose of all three drugs (Fig. S2, resulting in a dose dependency of interaction strength). We further show net (*DA*) interactions are antagonistic at a low dose and shift to additivity or synergy at a high dose (Fig. 4). Most of the dose-dependent transitions are from additivity (no interaction) to either synergy or antagonism. Transitions between synergy and antagonism—corresponding to an extremely abrupt or sharp transition—are extremely rare, at less than 4% for *DA* and less than 1% for *E3*. Antagonistic interactions remain antagonistic or transition to additivity more for 2-drug combinations (26%) than for 3-drug combinations (17%). Emergent interactions (*E3*) are rarely synergistic. No drug combinations exhibit emergent synergy at the low dose (index 1), while less than 4% do so at the high dose (index 6) (Fig. 4). Interaction transitions are summarized for each drug combination with both the sum and absolute change in *DA* (Fig. S3). Clearly, increasing the dose of one drug can lead to various trajectories for changes in *DA* (Fig. S4), such as a no change, additive to synergy, or additive to antagonism. Nevertheless, the landscapes are not randomly scattered with mixtures of interactions, but instead are composed of confined subspaces or regions of synergy or antagonism. Transitions between different interaction types are generally buffered by a region of additivity.

**Figure 4.**
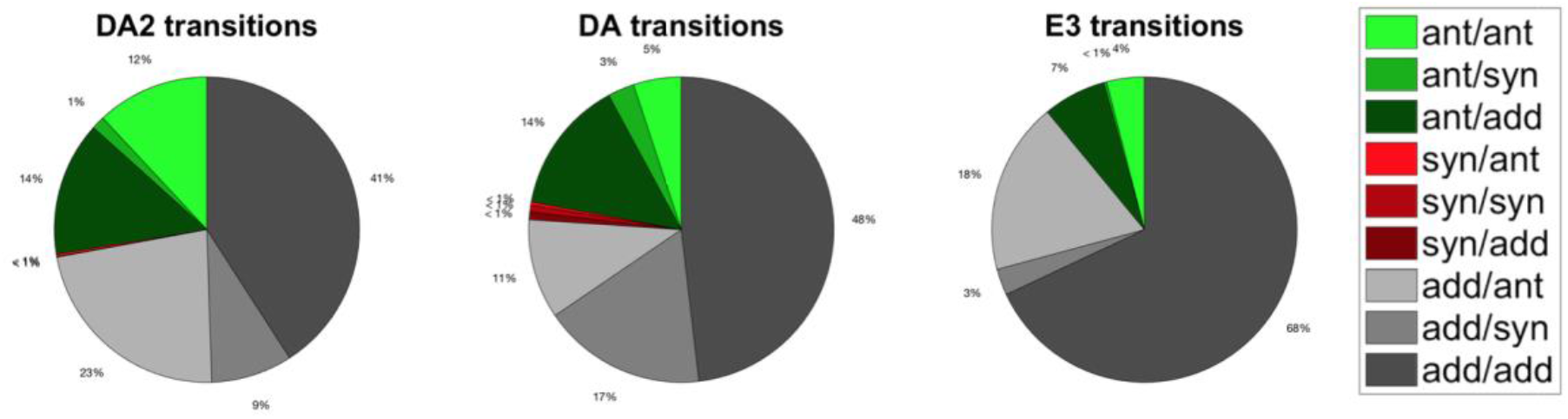
Distributions of transitions between interaction types from a low dose (index 1) to a high dose (index 6). To assess the effect of increasing dose on interaction in a two- drug case, we compared drug interaction of drug A at a sub-inhibitory concentration and drug B at either a high dose and a low dose. In a three-drug combination, the interaction was examined with a third drug at a high and low dose. Pairwise interactions (*DA2*, i.e. overall pairwise interaction) are dominated by antagonism and additivity at the low dose (green and gray, 99%), while a total of 10% are synergistic at the high dose (left). Three-way (*DA)* interactions are mostly additive at the low dose (gray, 76%) and antagonistic (green, 22%), but change from additivity to antagonism (16%) and from additivity or antagonism to synergy (21%) at the high dose. The emergent three-way interactions measured by *E3* are mainly additive at the low dose (gray, 89%) with the rest being antagonistic, and result in very few synergistic interactions at the high dose (3%), with some being antagonistic (22%) and a majority being additive (75%).

### Pairwise interactions contribute most to net three-way interactions, emergent three-way interactions are not predicted by pairwise interactions

For each triple-drug combination, we calculated three-way net (*DA*), pairwise net (*DA*), and three-way emergent (*E3*) interaction metrics for all possible drug pairs with a third drug at low, intermediate, and high concentrations (dose indices 2, 4, and 6). The relationship between the pairwise *DA* and three-way *DA* is mostly positive, with the net interactions strongly influenced by the pairwise interactions (Fig. S5). For all three doses, the mean of the three pairwise *DA* correlates strongly with three-way *DA* (Spearman’s *ρ* = 0.793, Fig. 5A). *DA* and *E3* show *no correlation* (Spearman’s *ρ* = –0.094, Fig. 5B), while *E3* and pairwise *DA* exhibit a slightly anti-correlated pattern (Spearman’s *ρ* = –0.384, Fig. 5C). A more synergistic mean pairwise *DA* thus predicts a more synergistic three-way *DA*, which is unsurprising, since the pairwise interactions are included in the three-way *DA*. Although the relationship between three-way *DA* and *E3* is weak or non-existent, the anti-correlation between mean pairwise *DA* and *E3* is striking. In particular, for synergistic three-way *DA* (red points) interactions, there is a strong anti-correlation between pairwise *DA* and *E3*, indicating that antagonistic pairwise interactions tend to be associated with strongly synergistic *E3*, which in turn drives the three-way interaction synergistic (Fig. S6). Conversely, when the mean of the three pairwise interactions is below zero (corresponding to synergy) and the three-way *DA* is synergistic, *E3* is predominantly antagonistic. This effect is likely to result from low fitness at high doses that can cause large deviations in *DA* and *E3*. The correlations between pairwise *DA*, three-way *DA*, and *E3* are similar for both low and intermediate doses of the third drug (dose indices 2 and 4), with similar correlation coefficients (Fig. S7).

**Figure 5.**
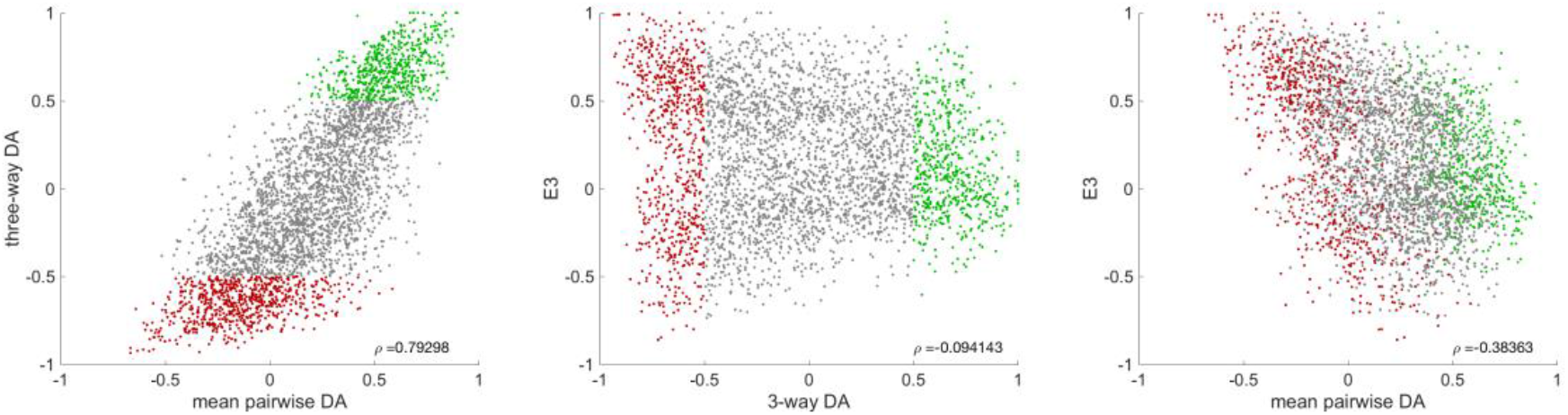
Comparisons of three-way interactions to pairwise interactions. For each three-drug combination, we calculated three-way net (*DA*), pairwise net (*DA*), and three-way emergent (*E3*) interaction metrics for all possible drug pairs and doses with the third drug at low, intermediate, and high concentrations (dose indices 2, 4, and 6). The relationship between the three-way *DA* at dose index 6 and the mean of the three pairwise *DAs* at the same doses shows a strong positive correlation (Spearman’s *ρ* = 0.793), whereas this correlation is absent between *E3* and three-way *DA*, as well as between *E3* and the mean pairwise *DA*. Together, this suggests that the *E3* interactions emerge independently of the pairwise interactions. The pairwise interactions surprisingly are negatively correlated with the emergent three-way interactions. *DA* and *E3* are evaluated at a high dose (dose index 6), and markers are colored according to the three-way *DA* value for antagonism (green), additivity (gray), and synergy (red). Three-way *DA* and *E3* are calculated for one drug at dose index 6 and the other two drugs at all dose combinations. The three pairwise *DA* components are calculated with one of the three drugs concentrations at zero. The mean pairwise-*E3* correlation for synergy alone is *ρ* = –0.619 (See Fig. S8).

## Discussion

To quantify the effect of dose on drug interactions, we measured fitness of a pathogenic strain of *E. coli* subjected to all possible 3-drug combinations of eight antibiotics across a gradient of doses for each drug. To visualize the high-dimensional interaction space of our data, we introduced the interaction landscape that displays quantitative measures of interactions as a function of the interacting components. We provide evidence that different environmental conditions (*i.e*., drug concentrations) can change drug interaction type and thus lead to dose-dependent interactions. We also showed these transitions are smooth, rarely going from synergy directly to antagonism or vice versa. Instead, transitions first pass through the intermediate type of additivity (no interaction) as they pass from antagonism to synergy or from synergy to antagonism.

The interaction landscapes give a direct and intuitive view of how the environmental space of combined drug doses affects the efficacy of drugs in combination. This representation is analogous to other maps of underlying control variables onto one dependent variable, such as genotype-fitness maps [21–23], genotype-phenotype maps [24, 25], and phenotype-fitness maps [26–28]. In addition, we expect our approach can be usefully applied to other systems related to toxins, pollution, and stressors.

Our results lead to two insights that should aid future studies of drug combinations. First, within our interaction landscapes, there are large, clearly delineated subspaces that correspond to specific types of drug interactions. These subspaces often occur at high or low concentrations of the combined drugs. Conclusions can therefore be made with less information than is needed for fitness landscapes by mapping the boundaries between these different subspaces and understanding how the magnitude of the interactions change when moving toward or away from a boundary. Moreover, these subspaces suggest simple methods for predicting regions of positive or negative evolutionary pressures on subpopulations of treated cells (*e.g*., selecting for or against resistance) and could have profound implications for choosing effective drug-dose combinations as well as intelligent drug treatments. Second, because there are transitions across the landscape that go between these subspaces of interaction type, synergistic combinations identified with only one dose regime [10, 13, 15, 29] can be antagonistic when used or prescribed at a different dose regime. Such a reversal could have detrimental impact on clinical decisions and scientific studies. For example, in figure 1C, an interaction type of antagonism at one set of doses ([ERY]= 125 μM, [AMP] = 0.39 μM, and [CLI] = 7.81 μM) changes to synergy at another set of doses ([ERY] = 125 μM, [AMP] = 6.25 μM, and [CLI] = 125 μM). Understanding how drugs interact in a dose-specific way will help to avoid conflicting results and potentially detrimental antagonistic combinations being applied in the wrong setting [30]. Importantly, fluctuating drug dosages could be used to create fluctuating selection pressures for cell populations. Indeed, evolutionary dynamics of a population can change drastically in changing environments [31, 32] and fluctuating environments can lead to higher levels of genetic diversity and biodiversity [33], evolution of generalist over specialist species [34], and other evolutionary and ecological phenomena. To assess whether this picture of drug interactions as strongly dose dependent goes beyond these particular drugs for this specific strain of *E. coli*, other drugs in other organisms need to be explicitly measured. Further detailed data and identification of general patterns across bacterial strains or drugs will contribute to better methods for predictions.

Zimmer *et al*. [15] proposed a model that predicts higher-order interactions at a full range of doses based only on pairwise interactions at low doses. We find component pairwise interactions are the largest contributor to overall net interactions which suggests the approach of Zimmer *et al*. may be frequently useful. However, pairwise interactions are independent of emergent interactions, so we doubt that higher order interactions will be easily predictable using the framework of Zimmer et al. That is, for most of the triple-drug combinations, the pairwise *DA* is a reasonable predictor of three-way net interaction (*DA*), but it does not correlate well with or usefully predict *E3*. Moreover, in some cases of synergistic three-way *DA* is not predicted by any component pairwise interactions, we do find a correlation between three-way *DA* and *E3*, showing the net interaction arises from the emergent part as would be expected. Our results are consistent with the basic findings of the Zimmer *et al*. model for net interactions, but show that emergent, higher-order interactions are independent and not predicted from component pairwise interactions. For cases where there are only three-way interactions, but very weak or non-existent pairwise interactions, inferences based on the Zimmer *et al*. model will therefore be especially misleading. This critical distinction seems absent from the literature because previous studies on dose-dependent interactions have been conducted with either limited numbers of drug combinations, with drugs at fixed doses, or have analyzed interactions with methods that do not distinguish net versus emergent interactions. Our study compensates for both the lack of data and missing analysis for emergent interactions for dose dependency.

Finally, we note that examining the whole-drug space for three-drug combinations can be extremely time consuming and expensive. An intriguing recent work by Cokol *et al. [35]* sampled data that correspond to a portion of our interaction landscape in order to infer the interaction type based on the Loewe additivity model in which it is assumed that a drug cannot interact with itself [36]. However, this methodology requires that the interactions be uniform throughout the entire interaction space, such that the contours stay either concave or convex across all doses. That is, the Cokol *et al*. framework assumes that there is no dose dependency, meaning no transitions between subspaces of interaction types. In contrast, our study using Bliss Independence models, which applies to single-dose measurements and makes no assumptions about dose dependency, shows that drug interactions generally are strongly affected by dose when we look at the entire interaction landscape. These fundamental discrepancies between the Bliss and Loewe models are also observed in two-drug interactions [37]. Future work to understand the meaning of these differences, which are intricately connected to the domain of Bliss versus Loewe models, are therefore greatly needed.

Our introduction of interaction landscapes along with our results that transitions are typically smooth and gradual should greatly aid in intuiting and thus understanding the complexity of drug interactions. Such insight is needed because combinatorial therapy is an extremely common practice in complex, chronic diseases such as hypertension, infectious disease, and cancer [38, 39], and could be strategically valuable for preventing the evolution of resistance. Visualization and analysis of multi-dimensional interaction data is a challenge faced by an increasing number of disciplines as experimental advances for collecting big data continue to grow. By combining our large dataset with a rigorous theoretical framework to quantify both net and emergent interactions, our approach enables new insights via the detection and quantification of how multi-drug interactions change with dose from low to high concentrations or for small or large numbers of drugs.

## Materials and Methods

### Bacterial Strain

We used *E. coli* CFT073, a highly virulent pyelonephritis strain isolated from human clinical specimen, obtained from ATCC (designation number 700928). The strain was grown in 2 mL of LB media (10 g/L tryptone, 5 g/L yeast extract, and 10 g/L NaCl) and streaked onto LB agar plates to isolate single colonies. Then a single colony was inoculated into 2 mL of LB and grown for 24 hours. Following the incubation, the culture was mixed with 2 mL of 50% glycerol and aliquoted into 50 μL to generate bacterial cell stocks with 25% glycerol for storing at –80°C. Each experiment was started with a thawed aliquot stock by inoculating 20 μL into 2 mL of LB media. The culture was incubated at 37°C until it reached exponential growth phase (an OD of 0.5) and diluted to maintain 10^4^ cells per experimental condition.

### Antibiotics

Antibiotics used include erythromycin (ERY), ampicillin (AMP), clindamycin (CLI), streptomycin (STR), nitrofurantoin (NTR), cefoxitin (FOX), and trimethoprim (TMP), all from Sigma (St Louis, Mo), and ciprofloxacin (CPR) from MP Biomedicals (Santa Ana, Ca). All antibiotics were dissolved and sonicated in 100% DMSO (Sigma) except for STR which was dissolved in 50% DMSO, due to limited solubility in 100% DMSO. Experiments for IC50 and drug interactions (below) were conducted in clear flat bottom 384-well plates from Greiner BioOne.

### IC50 determination

A 20-step two-fold serial dilution was prepared for each antibiotic. The source plate was made by preparing each drug with a total volume of 70 μL at 10 mM as the starting concentration, or the first step, filled into a 384 well plate. The following dilution steps were conducted by a robotic liquid handling system with a transfer volume of 35 μL per step. Meanwhile, 25 μL of LB per well were prefilled into a second 384-well plates using the Multidrop 384 (Thermo Scientific). Next, 500 nL from the source plate were delivered into the prefilled plate using the Biomek FX (Beckman Coulter) with a pin tool (V&P Scientific). Then, 25 μL of bacteria inoculum was added to each well to reach a final 50 μL per well with 1% DMSO. Each plate included negative controls (media alone), vehicle controls (media with 1% DMSO), and positive controls (media with 1% DMSO and cells). The plates were incubated at 37°C with OD_595_ measurement for cell density at 4-hour intervals for 24 hours. IC50s were determined by fitting a sigmoidal dose-response curve using the software Graphpad Prism.

#### Determining drug-dosage levels from dose-response curve of single drugs

To establish reasonable resolutions of various drug doses, we designed our dilution regime (Fig. 2A) to cover a wide range of dose effectiveness in terms of bacterial fitness of lethal, low, intermediate, and high. Mean dose response curves of each single drug (Fig. 2C) show a sigmoidal and monophasic curve that results in the desired fitness levels. Dose indices 1 and 2 are regarded as low doses, where fitness is between 1 and 0.8, with fitness here measured as growth rate relative to bacteria in no-drug environments. Dose indices 3 to 5 are intermediate doses that give a mean fitness around 0.4. High doses of 6 and 7 result in fitness below 0.2, except for clindamycin that has fitness well above the other drugs. We then calculated IC95 concentrations—where the dose concentration inhibits 95% of bacterial growth compared to no-drug environments—for each single drug (Table 1) to normalize the combined dose in triple-drug combinations in terms of combined effectiveness.

**Table 1.** List of drugs used in the study.

| Compound | Abbreviation | Cellular Target | Top dose ($\mu\text{M}$) | IC95 ($\mu\text{M}$) |
| --- | --- | --- | --- | --- |
| Ampicillin | AMP | cell wall synthesis inhibitor | 25 | 10.0 |
| Cefoxitin | FOX | cell wall synthesis inhibitor | 25 | 15.4 |
| Ciprofloxacin | CPR | Fluoroquinolone, DNA gyrase inhibitor | 0.5 | 0.1 |
| Nitrofurantoin | NTR | DNA damaging, multiple mechanisms | 250 | 56.7 |
| Trimethoprim | TMP | folic acid synthesis inhibitor | 10 | 4.8 |
| Streptomycin | STR | Aminoglycoside | 50 | 26.8 |
| Clindamycin | CLI | Protein Synthesis inhibitor, 50S | 500 | 722.3 |
| Erythromycin | ERY | Protein Synthesis inhibitor, 50S | 500 | 268.3 |

### Drug combination experiment

All three-drug combinations formed from a set of eight drugs were tested, resulting in 56 unique three-drug combinations. A source plate for each drug was prepared in seven-step, two-fold dilutions with various starting concentrations (Table 1), dependent on their respective IC50, with a total of 70 μL in DMSO at each dilution step. In addition, a zero dose was included into each drug gradient as the lowest concentration. A combination drug plate was prepared by pinning from each source plate of the component drugs using a 250 nL pin tool (V&P Scientific) to restrict the DMSO concentration to be lower than 1%. Methods for cell inoculation and incubation were the same as stated above. OD measurements were taken at 12 hours.

### Measuring fitness

Optical density measurements were made with Perkin Elmer Wallace 1420. Fitness was calculated as: 

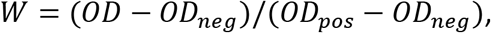

where *OD* is the optical density of the experimental condition with bacteria and drugs, *OD_pos_* is the positive control without drugs, and *OD_neg_* is the negative control without bacteria or drugs. Fitness is given with a precision of two decimals, and we therefore exclude fitness measurements below 0.01.

### Quantifying interactions

Interactions are commonly quantified as the deviation from Bliss independence [40]. We quantify this using *deviation from additivity* (DA), which measures interactions between drugs, while additivity is defined when the presence of one drug does not affect the percent reduction of bacterial growth of another drug. If the fitness of the organisms given three drugs is *W_XYZ_*, and the fitness when only given one drug is *W_X_, W_Y_, and W_Z_*, for drugs X, Y, and Z, respectively, then

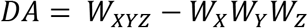

[41]. *DA* incorporates both pairwise and three-way drug interactions but cannot discern between them. To measure emergent interactions that occur only when three drugs are present, we subtracted both the single-drug effects and the component pairwise effects: 

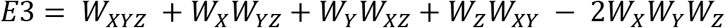

[41]. To delineate boundaries and tease apart interactions as synergistic, additive, and antagonistic from the unimodal distribution of *DA* and *E3*, rescaling was applied to each measurement. *DA* was rescaled by dividing by the absolute value of *DA*, but replacing *W_XYZ_* (denoted as 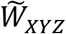) by 0 if *DA* ≤ 0, to account for cases of extreme lethal synergy (*W_XYZ_* = 0) while no single drug completely was completely lethal, and by the minimum value of *W_X_, W_Y_, W_Z_* if *DA* > 0, for cases of buffering antagonism where combined drugs have the same effect as the strongest single-drug effect. 

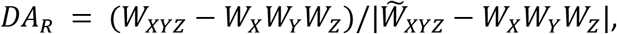

where 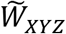 = 0 for *DA* ≤ 0, and min(W_X_, W_Y_, W_Z_) otherwise.

Similarly, *E3* is rescaled by dividing by 

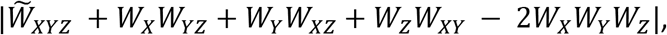

where 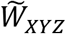 = 0 for *E3* ≤ 0 and min(W_X_W_YZ_, W_Y_W_XZ_, W_Z_W_XY_) otherwise [41]. This rescaling results in values between –1 and ∞. We further discretize both *DA_R_* and *E3_R_* based on the natural breaks in the histogram distribution of *DA_R_* and *E3_R_*. Values below –0.5 correspond to synergy, between –0.5 and 0.5 to additivity, and above 0.5 to antagonism. Values above 1 are capped at 1.

### Smoothing

To increase our confidence and resolution of interaction transition with dose, given the fairly noisy OD measurements, we smoothed the data using a weighted average algorithm by considering our dose combination matrix as a metric space. For each data point (interaction measurement at each drug-dose combination), both rescaled *DA* and *E3* were recalculated as a weighted average depending on the Euclidean distance (within the three-dimensional matrix) between the original data point and the points used for calculation. The weight is 1 for the origin, and 1/8*d* for the 26 nearest neighbors, where *d* is the Euclidean distance from the origin. If a neighboring value was missing, either because it lies at the boundary or because it was excluded due to low fitness, its weight was set to zero. The sum of the weights was required to comprise at least 59 percent of the original weight matrix. For smoothing, both *DA* and *E3* were truncated to values between –1 and 1, with higher values set equal to 1.

## Acknowledgements

We thank Nina Singh and Cynthia White for comments on the manuscript. We are grateful for funding from the Hellman Foundation (PJY), a KL2 Fellowship (PJY) through the NIH/National Center for Advancing Translational Science (NCATS) UCLA CTSI Grant Number where W UL1TR001881, and a James F. McDonnell Foundation Complex Systems Scholar Award (VMS). This investigation was also supported by National Institutes of Health, under Ruth L. Kirschstein National Research Service Award (T32-GM008185). Its contents are solely the responsibility of the authors and do not necessarily represent the official views of the NIH.

